# Simultaneous quantification of dynamic bacterial deformation and motility by machine learning

**DOI:** 10.64898/2026.07.07.737132

**Authors:** Kyosuke Takabe, Souichi Ugawa, Nobuo Koizumi, Shuichi Nakamura

## Abstract

We developed a convolutional neural network-based machine learning technique to simultaneously analyze the morphology and motility of spirochetal bacteria swimming with continuous cellular deformation. Matching probabilities between experimental images and learned models realizes quantification of cell morphology and association with motility. This method can be applied to diverse transformable cells, offering critical biophysical insights into microbial dynamics.

---

Bacterial motility exhibits remarkable diversity. Unlike species with extracellular flagella, such as *Escherichia coli*, spirochetes—including various pathogens responsible for severe infectious diseases—uniquely internalize their flagella within the periplasmic space beneath the outer membrane (Miyata et al., 2020; Nakamura, 2020). The rotation of these periplasmic flagella periodically deforms the cell body, thereby generating propulsive force (Nakamura, 2020). *Leptospira* spp., the focus of this study, possess a single periplasmic flagellum at each cell end, which transforms the cell extremities into either a “Hook” shape or a helical “Spiral” shape (Figure 1). The cell-end morphology frequently deforms, determining the swimming behavior: an asymmetric Spiral–Hook state drives the rotation of the protoplasmic cylinder to generate substantial propulsion, whereas symmetric configurations (Spiral– Spiral or Hook–Hook) counteract the generated torques and limit translocation (Goldstein and Charon, 1990; Nakamura et al., 2014; Takabe et al., 2017a,b) (See Supplementary Information for the detailed biological background of *Leptospira*).

**Figure 1.**
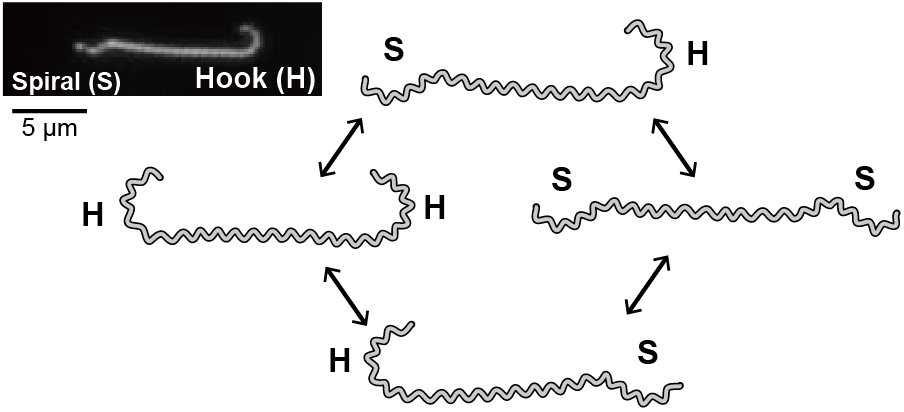
Morphological diversity of *Leptospira* cell ends. The left upper panel shows a dark-field micrograph of *L. biflexa*.

Despite this strong coupling between morphology and motility, the evaluation of these dynamic changes has long relied on manual, qualitative visual inspection (Goldstein and Charon, 1990; Takabe et al., 2017b), contrasting with well-established automated motility tracking techniques (Abe et al., 2023). To address this gap and eliminate human bias, this study introduces a machine learning-based approach to realize automated, quantitative measurement of *Leptospira* cell-end morphology and behavioral dynamics.

We employ a convolutional neural network (CNN) (Lecun et al., 1998) for our machine learning model implemented in PyTorch (Ansel et al., 2024), taking advantage of its well-established capability in two-dimensional image processing (Figure 2). The developed machine learning framework adopting CNN classifies the *Leptospira* morphologies into three distinct categories: Hook, Spiral, and Straight. While normal cell ends exhibit either Hook or Spiral, we assume that cell ends with defective flagellar expression, which are rarely observed, are recognized as Straight. Microscopic movie clips are first converted into 64 *×* 64–pixel grayscale images, followed by data augmentation using spatial transformations (flips, rotations, and transpositions) (Buslaev et al., 2020). To evaluate model performance, the dataset is randomly partitioned into training and validation sets using 5-fold cross-validation strategy. The CNN model is optimized using Stochastic Gradient Descent (SGD) with momentum to minimize the cross-entropy loss. For a batch size *N*, the loss function ℒ is defined as:

**Figure 2.**
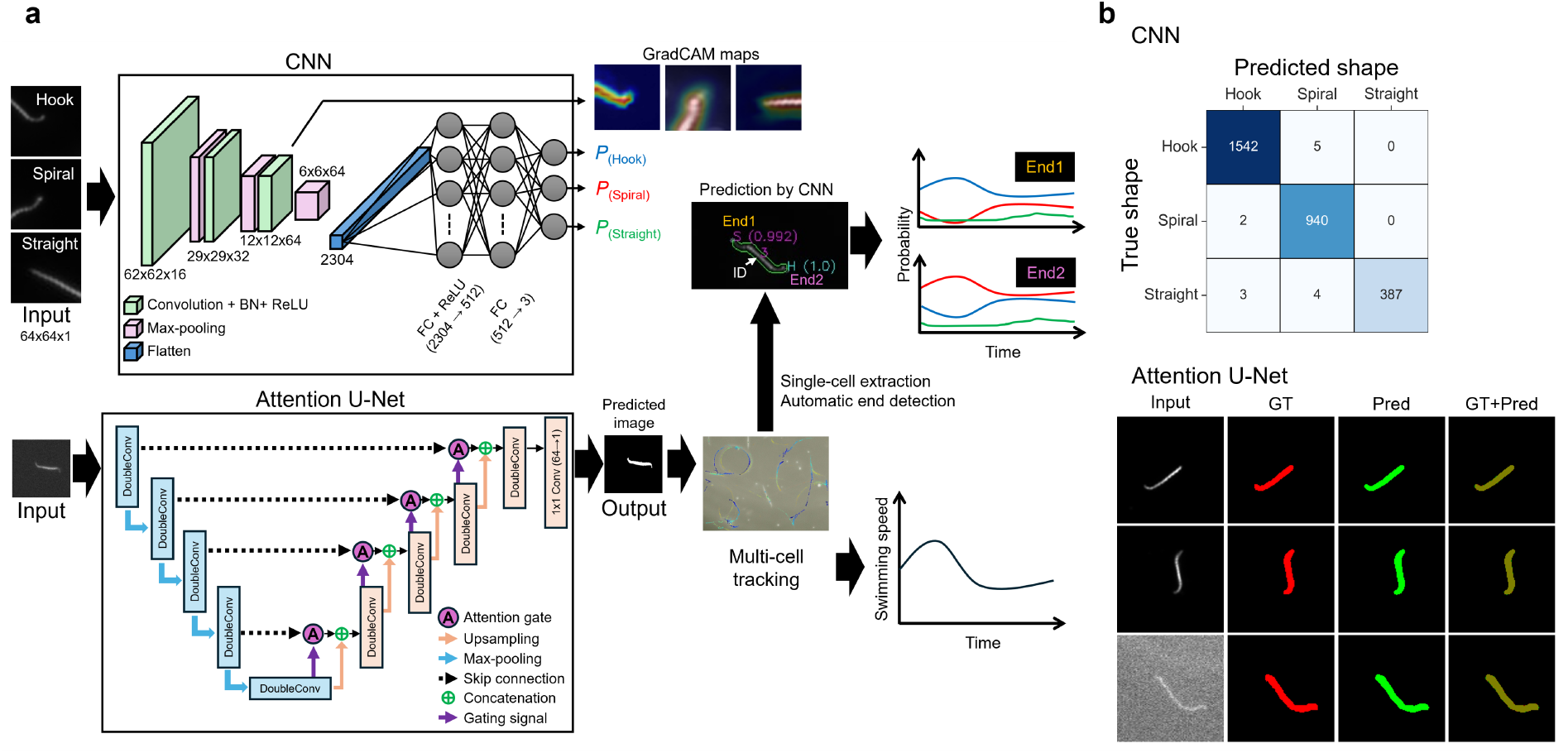
Automatic measurements of motility and morphological features. (a) Schematic diagram of our method, where a simple CNN for the end-shape classification and an Attention U-Net for single-cell detection are integrated. In the CNN architecture, each convolutional block (green) is composed of a Conv2d layer, batch normalization (BN), and a ReLU activation function, followed by two incorporated fully connected (FC) layers. For motility analysis, the predicted images from each frame are utilized to track multiple cells. Simultaneously, automatic end detection is applied to each tracked cell, allowing the CNN model to predict its end shape. Grad-CAM maps are generated using the gradients flowing into the final convolutional layer to verify the validity of the CNN model. Finally, single-cell time-course data for both motility and morphological parameters are obtained. (b) Model evaluation. For the CNN, the cumulative confusion matrix from 5-fold cross-validation demonstrates the high predictive performance of our model. For the Attention U-Net model, the visual comparison between ground-truth (GT) and predicted images (Pred), where each row corresponds to an individual cell, indicates the high precision of the model.

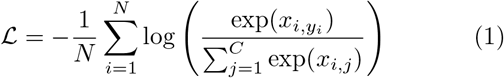

where *x*_*i,j*_ represents the model output (logit) of the *i*-th sample for class *j, y*_*i*_ is the ground-truth label, and *C* denotes the total number of classes (*C* = 3). To prevent overfitting, an early stopping mechanism monitors the validation loss during training. The learning curves for training and validation indicated that the model converged without overfitting (Figure S1a). A total of 2,933 images, comprising 1,570 for Hook, 961 for Spiral, and 402 for Straight, were used for learning. Consequently, the average accuracy reached 0.995 ± 0.005, and other metrics—precision, recall, and F1-score—were above 0.99, ex-cept for the recall of Straight (0.98) (Figure S1a). Cumulative confusion matrix from 5-fold cross-validation showed the high classification performance of our model, despite the biased training dataset (Figure 2b).

Utilizing the trained models, an automated cell-tracking pipeline is implemented, correlating bacterial motility with real-time morphological dynamics. The segmentation model (based on Attention U-Net architecture (Oktay et al., 2018)) was trained using over 14,000 annotated images with diverse brightness distributions. The model was optimized using Adaptive Moment Estimation (Adam) and achieved an average Dice score of over 0.97 from 5-fold cross-validation incorporating early stopping to monitor validation loss (Figure S1b). Representative results showed a close match between the ground-truth and predicted images (Figure 2b), indicating that the model can accurately segment individual cell regions with high precision. The prediction of cell regions by the segmentation model is followed by tracking individual bacteria across consecutive frames using a nearest neighbor displacement strategy bounded by a maximum relocation radius. Gradient-weighted Class Activation Mapping (GradCAM (Gildenblat and contributors, 2021; Selvaraju et al., 2019)) is integrated to compute visual explanations of the model’s predictions, tracking the localized morphological features that contribute to the classification.

The developed analysis framework was applied to evaluate the morphological and behavioral dynamics of *Leptospira biflexa* strain Patoc I, a canonical non-pathogenic spirochete (see Supplementary Information for details on the experimental procedure). Figure 3a displays time courses of the cell-end shape, morphological asymmetry, and displacement of a single *L. biflexa* cell. The cell ends were arbitrarily designated as End1 and End2. Cells with a straight end were rarely observed and were therefore excluded from the current analysis. Let *P*_1H_ and *P*_1S_ denote the probabilities that End 1 is in the Hook and Spiral states, respectively. Similarly, *P*_2H_ and *P*_2S_ represent the corresponding probabilities for End 2. We define the asymmetry index *α* as follows:

**Figure 3.**
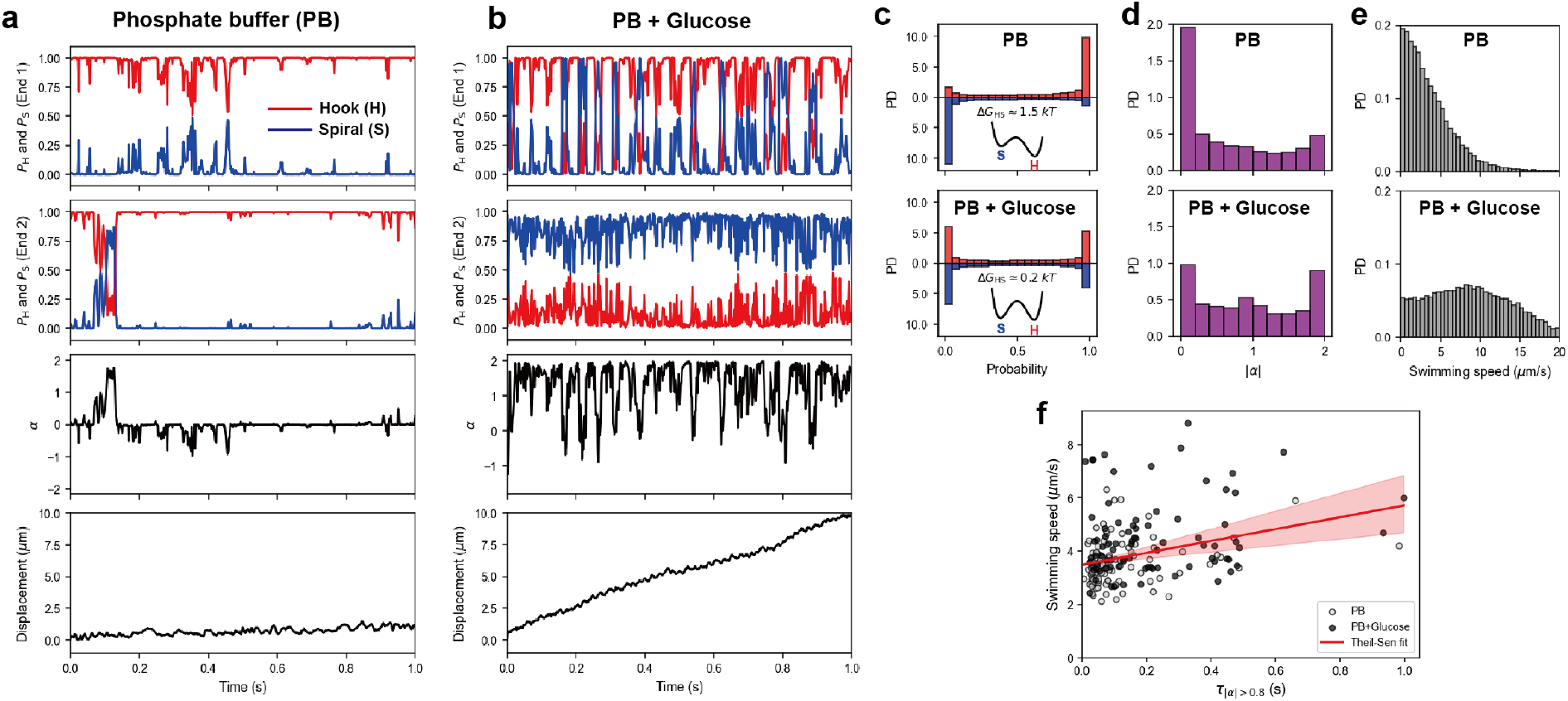
Morphological and motility analyses based on the CNN model. (a) Time courses of a leptospiral cell swimming in sodium phosphate buffer (PB, pH 7.4) and (b) in PB supplemented with 100 mM glucose (PB + Glucose), a chemotactic attractant for *Leptospira* spp. (Islam et al., 2014; Lambert et al., 2012). The upper two panels show the shape probabilities of the hook (red) and spiral (blue) forms for end 1 and end 2, respectively. The morphological asymmetry index *α* is calculated from these shape probabilities to quantitatively evaluate the degree of asymmetry (see main text). The bottom panels show the time courses of bacterial displacement. (c–e) Probability density (PD) distributions of (c) *P*_H_ (red) and *P*_S_ (blue), (d) |*α*|, and (e) swimming speed in PB (upper; 3.58 *±* 2.95 *µ*m/s) and PB + Glucose (lower; 8.63 *±* 4.97 *µ*m/s). Data from end 1 and end 2 are combined in these analyses. In (c), the free-energy difference between the hook and spiral states (Δ*G*_HS_) is calculated from the ratio of their appearance fractions ([S] and [H]) based on the thermodynamic relation [S]*/*[H] = exp(*™*Δ*G*_HS_*/k*_B_*T*). The values of Δ*G*_HS_ and the corresponding schematic energy landscapes are shown in the insets. (f) Relationship between the duration of the asymmetric morphology (*τ*_|*α*|*>*0.8_) and swimming speed. The Spearman’s rank correlation coefficient is *ρ* = 0.315 (*P* = 1.61 *×* 10^*™*5^). The solid red line represents the fit using the Theil–Sen estimator, and the shaded region indicates the 95% confidence interval. In (c–f), the number of analyzed cells is *n* = 33 for PB and *n* = 46 for PB + Glucose.

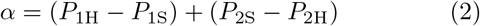

A higher absolute value of *α* indicates a more pronounced structural mismatch between the two ends of the cell. In phosphate buffer, the cell-end morphologies were strongly biased to the Hook shape, resulting in the predominant symmetrical morphology (*α* ~ 0) and suppression of net displacement, consistent with our previous study (Takabe et al., 2017b). The addition of glucose enhanced the end-shape variation and emergent asymmetric configuration (*α* ~ 2 at most), enabling directed bacterial traveling (Figure 3b). Figures 3c–f summarize the collective data from multiple cells based on these individual cell measurements. Reflecting the result that the end morphology was strongly biased toward the hook shape in the absence of glucose, the free-energy difference between the spiral and hook states (Δ*G*_HS_) was approximately 1.5 *k*_B_*T*; the addition of glucose then switched the thermodynamic stability of the two states to a comparable level (Figure 3c). Furthermore, the addition of glucose induced a population-level shift from symmetrical to asymmetrical cell morphology (Figure 3d) and accelerated swimming (Figure 3e). These findings demonstrate a positive correlation between the duration of the asymmetric morphology (*τ*_|*α*|*>*0.8_) and the swimming speed (Figure 3f).

In conclusion, the CNN-based image analysis method developed in this study successfully quantified the dynamically changing cell morphology of *Leptospira*. This capability enabled the thermodynamic characterization of morphological transitions and allowed us to elucidate their correlation with cell motility through our newly defined asymmetry index. Furthermore, the application of this approach extends beyond *Leptospira*; it holds great promise for analyzing other cell types where deformation plays a pivotal role in function, such as amoebae and leukocytes. This method will broadly contribute to the advancement of quantitative research across the life sciences.

## Fundings

This work was supported by the JSPS KAKENHI: 24K18444 for KT, 22K07062 for NK, and 22H04828 and 24K02274 for SN.

## Data Availability

The data supporting the findings of this study are available from the corresponding author upon request.

## Code Availability

The computer codes used for this study are available from the corresponding author upon reasonable request.

## CRediT authorship contribution statement

**Kyosuke Takabe:** Writing – original draft, Writing – review & editing, Software, Methodology, Investigation, Validation, Formal analysis, Data curation, Conceptualization, Funding acquisition. **Soichi Ugawa:** Investigation, Visualization, Validation. **Nobuo Koizumi:** Writing – review & editing, Resources, Funding acquisition, Conceptualization. **Shuichi Nakamura:** Writing – original draft, Writing – review & editing, Investigation, Visualization, Methodology, Validation, Formal analysis, Data curation, Conceptualization, Funding acquisition, Project administration, Supervision.

## Competing interests

The authors declare no competing interests.

## Declaration of generative AI and AI-assisted technologies in the manuscript preparation process

During the preparation of this work the authors used ChatGPT 5.5 (OpenAI) and Gemini 3.5 Flash (Google) as supplementary tools to assist with code development, including debugging and optimization. After using these tools, the authors reviewed and edited the content as needed and take full responsibility for the content of the published article.

## Notes

### Competing Interest Statement

The authors have declared no competing interest.

