## Supplementary Information for "Simultaneous quantification of dynamic bacterial deformation and motility by machine learning"

Title: Quantifying dynamic bacterial deformation and motility simultaneously via machine learning

### Evaluation of the CNN and the Attention U-Net models

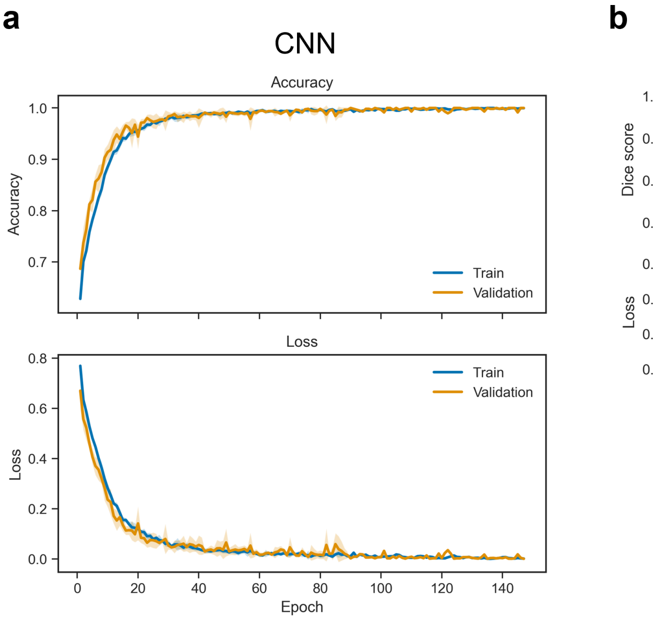


| Class | Precision | Recall | F1-score |
|---|---|---|---|
| Hook | 0.997 ± 0.004 | 0.997 ± 0.003 | 0.997 ± 0.003 |
| Spiral | 0.991 ± 0.011 | 0.998 ± 0.005 | 0.994 ± 0.008 |
| Straight | 1 | 0.982 ± 0.026 | 0.991 ± 0.014 |

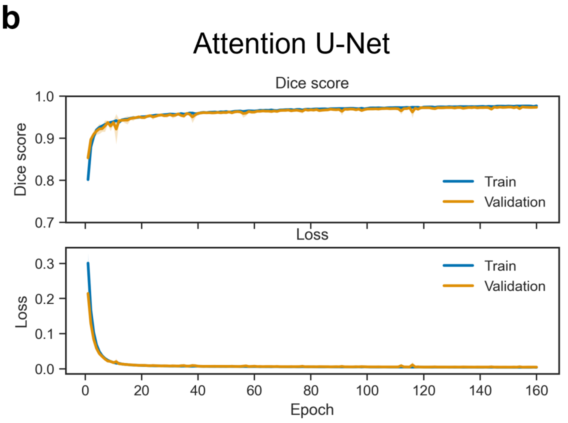


| Dice score | IoU | Precision | Recall |
|---|---|---|---|
| 0.972 ± 0.002 | 0.945 ± 0.003 | 0.973 ± 0.002 | 0.970 ± 0.003 |

Figure S 1. 5-fold cross-validation results for the two models. (a) Training and validation accuracy and loss curves of the CNN model as a function of epoch, with the standard deviation for each epoch presented as a shaded area, indicating that the model converged without overfitting. The table below presents the evaluation metrics for each class across the 5 folds. Values are presented as mean ± standard deviation. (b) Training and validation Dice score and loss curves of the Attention U-Net model as a function of epoch. The table below presents the evaluation metrics for the segmentation task.

### Detailed biological and biophysical backgrounds of *Leptospira*

Bacterial motility mechanisms vary widely across phyla. Well-studied model species such as *Escherichia coli* swim via the rotation of extracellular helical flagellar bundles, while other wall-less bacteria like *Mycoplasma* spp. utilize a specialized leg-like machinery on the cell surface to glide on solid substrata [1, 2]. In contrast, the phylum Spirochaetes is characterized by an elongated spiral or wavy cell body with internal periplasmic flagella. This phylum includes major human pathogens responsible for severe infectious diseases, such as *Treponema pallidum* (syphilis) and *Borrelia burgdorferi* (Lyme disease) [3].

The genus *Leptospira* encompasses both saprophytic and pathogenic species; the latter are the causative agents of leptospirosis, a re-emerging zoonosis that frequently breaks out in tropical and subtropical regions [4]. For these pathogenic spirochetes, active motility is a crucial virulence factor required for tissue invasion and dissemination, making the elucidation of their swimming mechanism a potential key to developing novel preventive and therapeutic strategies.

The bidirectional translocation of *Leptospira* is driven by the gyration of its two cell ends. The intracellular flagellum at each end is curved by asymmetrically localized flagellar sheath proteins [5], transforming the cell extremities into either a Hook shape or a helical Spiral shape (Figure 1). When viewed from the distal end of the cell, the Hook end gyrates counterclockwise (CCW), whereas the Spiral end gyrates clockwise (CW) [6, 7]. Although the direct contribution of Hook-end gyration to propulsion is minor, its counter-torque drives the CW rotation of the right-handed helical protoplasmic cylinder (PC, pitch size $\approx 0.6\,\mu$m) across the entire cell body, generating a substantial propulsive force. When the cell morphology is symmetric (Spiral–Spiral or Hook–Hook), the torques generated at both ends counteract each other, suppressing PC rotation and limiting apparent translocation. Consequently, increasing the probability of entering the asymmetric Spiral–Hook state directly enhances the migration capacity and environment-exploration efficiency of *Leptospira* [6–9].

### Biological sample preparation and experimental procedure

*L. biflexa* cells were cultured in EMJH medium at 30 °C until the late-logarithmic growth phase, diluted 1:10 with 20 mM phosphate buffer (pH 7.4), and introduced into a flow chamber constructed using slide and coverslips sealed with double-sided tape. Bacterial movement was observed under a dark-field microscope (BX50, Olympus, Japan) equipped with a 40× objective (UPlan FL N) and an oil-immersion dark-field condenser. Sequential images were recorded using a high-speed CMOS camera (ORCA-Fusion, Hamamatsu Photonics, Japan) at 500 frames per second ($\Delta t = 2$ ms) and analyzed using our machine learning pipeline (Figure 2).

Parameters characterizing the bacterial morphology and motility are determined at each time point (frame) for individual cells as follows. Morphological features include the orientation angle ($\theta$) obtained via ellipse fitting, the geometric centroid coordinates ($x_c, y_c$), the instantaneous swimming speed $v(t) = \sqrt{(\Delta x_c)^2 + (\Delta y_c)^2}/\Delta t$, and the posterior probabilities of the three distinct cell extremity shapes predicted by the CNN model ($P_{\mathrm{Hook}}$, $P_{\mathrm{Spiral}}$, and $P_{\mathrm{Straight}}$), which naturally satisfy the conservation relation $P_{\mathrm{Hook}} + P_{\mathrm{Spiral}} + P_{\mathrm{Straight}} = 1$.
